# A molecular toolkit for the green seaweed *Ulva mutabilis*

**DOI:** 10.1101/2020.12.15.422947

**Authors:** Jonas Blomme, Xiaojie Liu, Thomas B. Jacobs, Olivier De Clerck

## Abstract

The green seaweed *Ulva* is an ecologically-important marine primary producer as well as a promising cash crop cultivated for multiple uses. Despite its importance, several molecular tools are still needed to better understand seaweed biology. Here, we report the development of a flexible and modular molecular cloning toolkit for the green seaweed *Ulva mutabilis* based on a Golden Gate cloning system. The toolkit presently contains 125 entry vectors, 26 destination vectors and 107 functionally validated expression vectors. We demonstrate the importance of endogenous regulatory sequences for transgene expression and characterize three endogenous promoters suitable to drive transgene expression. We describe two vector architectures to express transgenes via two expression cassettes or a bicistronic approach. The majority of selected transformants (50-80%) consistently give clear visual transgene expression. Furthermore, we made different marker lines for intracellular compartments after evaluating 13 transit peptides and 11 tagged endogenous *Ulva* genes. Our molecular toolkit enables the study of *Ulva* gain-of-function lines and paves the way for gene characterization and large-scale functional genomics studies in a green seaweed.

**One-sentence summary:** Molecular cloning tools allow to generate gain-of-function seaweed lines that will help to study seaweed biology.

## Introduction

Important progress in molecular research has been made for several unicellular algae, including comparative genomics (Blaby-Haas and Merchant, 2019), genetic transformation (*e.g*. (Doron et al., 2016; Faktorová et al., 2020)) and large systems-biology studies (*e.g*. (Mackinder et al., 2017; Strenkert et al., 2019)). Multicellular algae, or seaweeds, are important organisms in different marine habitats and several species are cultivated for food, feed, biofuel or pharmaceuticals (Bolton et al., 2016; Charrier et al., 2017; Wells et al., 2017). Notwithstanding their importance, molecular tools are needed to study seaweed biology and generate new strains with desirable characteristics (Loureiro et al., 2015). While genomic resources are increasing for brown, green and red seaweeds (Cock et al., 2010; Collén et al., 2013; De Clerck et al., 2018; Wang et al., 2020), stable genomic integration and expression of exogenous DNA is still rare and limited to antibiotic resistance genes and/or reporter genes such as *lacZ* or *GFP (e.g*. (Jiang et al., 2003; Song et al., 2003; Son et al., 2012; Lim et al., 2013; Uji et al., 2014; Oertel et al., 2015; Lim et al., 2019)). Recently, an attempt to overexpress a *Pyropia yezoensis* gene coding for a light harvesting protein resulted in co-suppression (Zheng et al., 2020). To our knowledge, no transgenic macroalgal line has been described expressing an endogenous tagged gene. Developing molecular tools for macroalgae will allow the study of gene function, which is crucial to understand seaweed growth, reproduction and dependence on symbiotic bacteria (Charrier et al., 2017).

The green seaweed *Ulva* is a multicellular marine algae model that has traditionally been studied from a morphological and physiological perspective (reviewed in (Wichard et al., 2015)), but is also a relevant commercial species (Bolton et al., 2016). Most research has been conducted on *U. mutabilis*, a species that has been studied and kept in culture since the 1950s (Føyn, 1958; Wichard et al., 2015). Advantages of this model organism are the short life cycle, ease of cultivation in well-defined media and the ability to generate stable transformants using plasmid vectors (Stratmann et al., 1996; Spoerner et al., 2012; Oertel et al., 2015). Importantly, the complete *U. mutabilis* genome is published and annotated (De Clerck et al., 2018). Transcriptomes, organelle genomes or specific loci have been reported in other *Ulva* species (Melton III et al., 2015; Zhou et al., 2016a; Zhou et al., 2016b; Cai et al., 2017; Yamazaki et al., 2017; Ichihara et al., 2019; Wang et al., 2019). Despite the availability of genomic resources, few molecular tools are described for investigating gene function in *Ulva*. So far, only four *Ulva*-specific promoter sequences have been isolated (Kakinuma et al., 2009; Oertel et al., 2015; Wu et al., 2017) and the expression of three transgenes has been reported (*bleomycin resistance* (*ble*), *GFP* and *lacZ;* (Huang et al., 1996; Kakinuma et al., 2009; Suzuki et al., 2014; Oertel et al., 2015)). Stable *U. mutabilis* transformants can be generated through expression of a *ble* cassette containing the promoter, first intron and terminator of Rubisco *SSU* (denoted here as BleR; (Oertel et al., 2015)).

Efficient modular cloning systems are a prerequisite in the design of large-scale functional genomic studies and synthetic biology. Flexible molecular toolkits based on Golden Gate cloning allow efficient assembly of expression cassettes (Lampropoulos et al., 2013; Casini et al., 2015). For example, a modular library containing 119 parts has been developed for the unicellular green algae *Chlamydomonas reinhardtti*, including seven promoters, 11 signal and targeting peptides, 34 reporter genes and eight antibiotic resistance genes (Crozet et al., 2018). In general, new modules are easily made and their modularity allows the generation and testing of a multitude of vectors in a straightforward way. At present, no such system is available for seaweeds.

In this study, we present a modular vector toolkit for molecular research in *U. mutabilis* based on the GreenGate cloning system (Lampropoulos et al., 2013). The toolkit presently contains 125 entry vectors, 26 destination vectors and 107 expression vectors. We demonstrate the importance of using endogenous promoter and intron regions for proper transgene expression and report stable overexpression of different transgenes using microscopy and molecular methods. The toolkit allows the rapid generation of transgenic *Ulva* lines expressing tagged transgenes, enabling the study of gain-of-function lines in green seaweeds.

## Results

### A universal cloning method and repository for *Ulva* vectors

An important prerequisite for a molecular toolkit is a universal and cost-effective cloning method that can be easily adopted by different laboratories. To this end, we generated a set of plasmids based on the GreenGate cloning system ((Lampropoulos et al., 2013); Fig. 1a; Table S1). This Golden Gate cloning system allows rapid and efficient assembly of up to six building blocks in a single-tube reaction. Every building block, or entry vector, is flanked by convergent BsaI restriction enzyme recognition sites. Upon BsaI cleavage, pre-defined 4-nt 5’ overhangs are generated on both ends of the DNA fragment that ensure the building blocks are assembled in the desired order and orientation in the destination vector (Lampropoulos et al., 2013). This modular assembly method is both fast (see Materials & Methods) and inexpensive. The main disadvantage is the need to ‘domesticate’ all sequences by removing internal BsaI sites. When building blocks are made, vector assembly, *E. coli* transformation and plasmid quality checks with restriction digest and Sanger sequencing can be done in as little as four days. After assembly and quality control steps have been met, 5μg of plasmid DNA is used to transform *Ulva* gametes ((Oertel et al., 2015); Fig. 1b). *Ulva* gametes are haploid and develop parthenogenetically after settling. Transformants are selected using phleomycin, undergo normal development with the help of at least two bacterial symbionts (*Roseobacter* MS2 and *Cythophaga* MS6; (Spoerner et al., 2012)) and are fully grown after ~4-5 weeks in Ulva Culture Medium (UCM; Fig. 1b). In summary, new vector combinations can be designed and tested in *Ulva* stable transformants in 5-6 weeks. All vectors generated in this study (Table S1) are available through https://gatewayvectors.vib.be.

**Figure 1.**
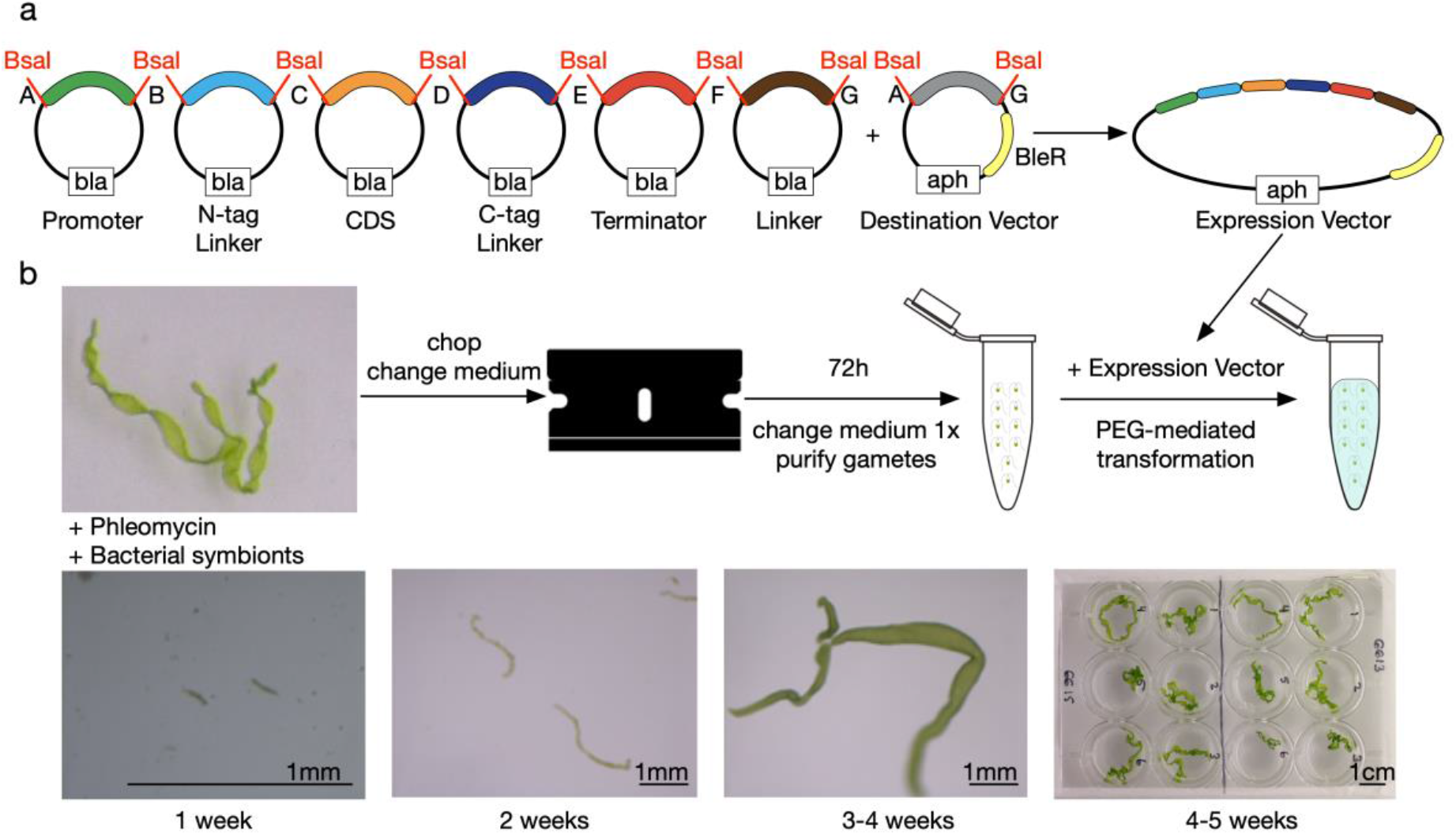
Overview of cloning strategy and transformation of *Ulva mutabilis*. a) Summary of GreenGate cloning strategy (Lampropoulos et al., 2013). Up to six entry modules (depicted with A-B, B-C, C-D, D-E, E-F and F-G) can be assembled in one destination vector in one reaction generating an expression vector. All entry modules have an ampicillin/carbenicillin (bla) resistance cassette for bacterial selection, the destination vectors contain a kanamycin (aph) resistance cassette. Although any sequence can be cloned in all entry modules, the most common sequences in every module are depicted (*e.g*. promoter sequence in A-B module). BleR is a module containing the ble gene, encoding bleomycin resistance protein, adapted for *Ulva* expression (see (Oertel et al., 2015)). b) Overview of *Ulva* transformation and selection. Gametogenesis of mature *Ulva* is induced by fragmentation and changing medium to wash out sporulation inhibitors. After 72h, the cells have differentiated into gametes that are released from the tissue after one more medium change. Gametes are isolated using their phototactic properties and subsequently DNA can be transfected using PEG-mediated transformation (Oertel et al., 2015). Transformed gametes develop parthenogenically into new thalli and can be selected using phleomycin. Representative images of transformants at different developmental stages.

### Endogenous promoters ensure transgene expression in *Ulva*

The promoter region of the small subunit of Rubisco (*pRbcS2*) is the only promoter functionally validated in *Ulva* by expression of BleR, (Oertel et al., 2015). Therefore, we designed a promoter screen to expand the number of functional promoters for *Ulva*. We first identified 43 highly expressed genes from previously generated transcriptome data (De Clerck et al., 2018) and selected genes that had a consistently high expression value greater than 3,000 FPKM. We narrowed this list down to 14 genes (Fig. 2a; Fig. S1a; Table S1) by removing those with internal BsaI and BbsI restriction sites in the promoter regions. These genes mainly code for ribosomal proteins or proteins involved in photosynthesis (Fig. S1a).

**Figure 2.**
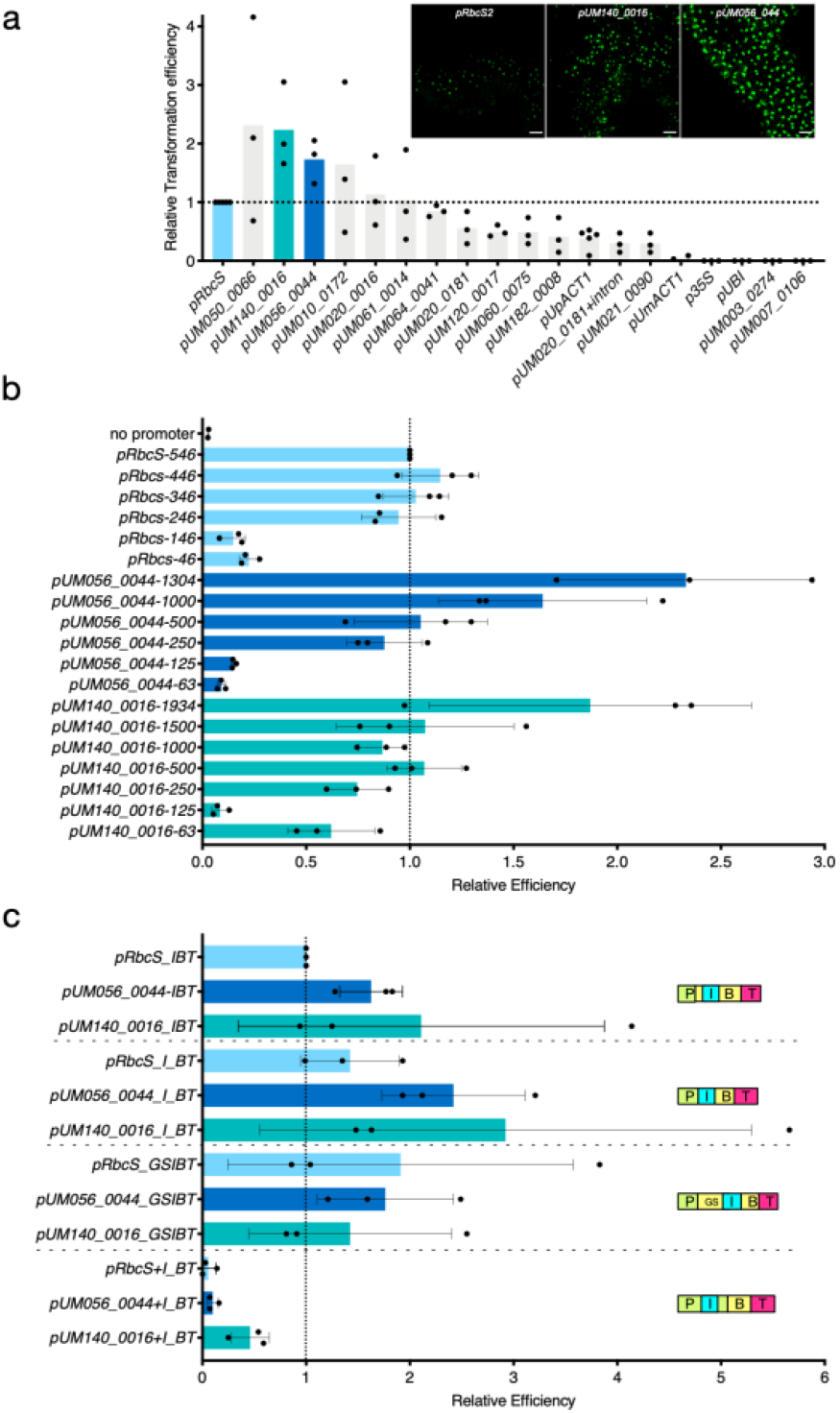
Functional validation of *Ulva* promoter and intron sequences. a) Twenty different regulatory promoter regions were cloned upstream of the resistance cassette (BleR-GFP; (Oertel et al., 2015)). To evaluate the relative strength of each promoter, the number of resistant individuals was counted relative to the number of resistant individuals counted for pRbcS ((Oertel et al., 2015); Figure S1). Visualisation of BleR-GFP expression is included for constructs containing *pRbcS, pUM140_0016* and *pUM056_0044* (See Fig. S2b for other lines). Each dot represents relative transformation efficiency for one independent experiment, the mean is indicated with a bar. n=3-5. b) Promoter deletion experiment for three promoters (*pRbcS, pUM140_0016* and *pUM056_0044*). The size of the promoter region is indicated for each independent line. Resistant individuals were counted and relative transformation efficiency was calculated as in (A). A control without promoter sequence was included. Each dot represents relative transformation efficiency for one independent experiment, the mean is indicated with a bar, error bars represent standard deviation. n=3. c) Importance of intron sequences for transgene expression in *Ulva*. The *Rbcs2* intron is originally an integral part of the BleR cassette (IBT; (Oertel et al., 2015)). The *Rbcs2* intron was cloned as a separate module directly upstream of the translation start site (I_BT), incorporated in the BleR gene with a G4SGS protein linker (GSIBT) or incorporated in one of three promoter regions (*pRbcS+I*; *pUM140_0016+I* or *pUM056_0044+I*). The number of resistant individuals was counted and transformation efficiency was calculated relative to pRbcS_IBT. Each dot represents relative transformation efficiency for one independent experiment, the mean is indicated with a bar, error bars represent standard deviation. n=3.

We confirmed the expression of the selected genes in *Ulva* individuals using qRT-PCR (Fig. S2a) and cloned the 770bp-2081bp region upstream of the predicted translation start site (TSS). Additionally, we generated one clone that contained the promoter region of the gene (*UM020_0181*) and its putative leader intron (Fig. S1a). We assembled the promoters to the *BleR* resistance gene directly fused to *Green Fluorescent Protein* (*GFP*; (Fuhrmann et al., 1999)) and the terminator of *RbcS2* (*promoter-BleR-GFP-tRbcS*; Fig. 2a; Fig. S1b). Similarly, we generated *pRbcS-BleR-GFP-tRBCS* as a positive control (546bp; (Oertel et al., 2015)) and generated expression vectors with the promoter region of an endogenous actin gene (*pUmAct1*; 2kbp), the previously characterized *U. prolifera* actin1 promoter (*pUpAct1*; 1941bp; (Wu et al., 2017)), the Cauliflower Mosaic Virus 35S promoter (*p35S*; 864bp; (Odell et al., 1985)) and the Arabidopsis *UBIQUITIN10* promoter (*pUBI*; 633bp; (Norris et al., 1993)). *Ulva* gametes were transformed with the expression vectors and the number of resistant individuals were counted for each promoter variant. To assess the relative strength of the promoters, we counted the number of resistant individuals and compared this to the number of resistant individuals generated using the positive control *pRbcS* ((Oertel et al., 2015); Fig. S1c). Generally, around 100 total transformants were obtained for control *pRbcS* transformations. Fourteen promoters resulted in phleomycin-resistant individuals (Fig. 2a) whereas five promoter regions did not lead to any resistant individuals (*pUM003_0274, pUM007_0106, pUmAct1, p35S* and *pUBI*). Two promoters (*pUM140_0016* and *pUM056 0044*) resulted in ~two-fold more resistant individuals compared to *pRbcS*, five promoters were more variable or similar to *pRbcS (pUM010_0172, pUM020_0016, pUM064_0041, pUM050_0066* and *pUM061_0014*) and six promoters generated fewer events than *pRbcS (pUpACT1, pUM120_0090, pUM021_0090, pUM182_0008, pUM060_0075* and *pUM020_0181*). Inclusion of a putative leader intron with *pUM020_0181* did not increase the strength of this promoter (Fig. 2a).

To qualitatively evaluate transgene expression in mature transformants, we imaged the BleR-GFP protein using confocal microscopy. We were able to visualize nuclear GFP expression for each functional *promoter-BleR-GFP-tRbcS* combination in mature thalli of transformants (Fig. 2a; Fig. S2b).

In addition to the promoter screen, we assessed the effect of successive deletions on the functionality of *pRbcS* and our two strong promoters *pUM140_0016* and *pUM056_0044*. *pRbcS* is a short sequence of only 546bp, so we successively deleted 100bp. For *pUM140_0016* and *pUM056_0044* the size of the promoter was successively halved (1000bp-500bp-250bp-125bp-63bp; Fig. 2b). Similar to our promoter screen, we transformed *Ulva* gametes with all constructs, counted number of resistant individuals and represent the strength of the constructs relative to the control *pRbcS* (Oertel et al., 2015). In general, we observed that for all three promoters the successive deletion of fragments negatively affects the relative transformation efficiency. Promoter sequences ≥250bp confer a comparable or higher relative transformation efficiency compared to *pRbcS* (Fig. 2b). When promoter size is smaller than 250bp, the number of resistant individuals drops sharply, indicating that the main transcriptional enhancers are located in this region (Fig. 2b). One intriguing exception is the 63 bp promoter of *pUM140_0016*, which shows an average relative transformation efficiency of 0.62; whereas this value is 0.08 for the 125 bp promoter of this construct (Fig. 2b).

### Endogenous introns ensure transgene expression in *Ulva*

Heterologous genes introduced in green algal genomes are often poorly expressed. For example, only 13% of *Chlamydomonas* transformants with a simple *VENUS*-containing plasmid (pMO518) show fluorescence (Onishi and Pringle, 2016). One way to increase transgene expression is to incorporate endogenous regulatory sequences in the coding sequence (Lumbreras et al., 1998; Oertel et al., 2015; Jaeger et al., 2019). To evaluate the importance of the endogenous *Ulva RbcS2* intron (*RbcsI*) sequence incorporated in the BleR coding sequence (Oertel et al., 2015), we generated intron variants of the resistance cassette. We incorporated the *RbcsI* sequence either as a stand-alone sequence before the TSS of BleR, integrated in a sequence encoding a small G4SGS protein linker directly upstream of the coding sequence of BleR or incorporated in one of three promoter sequences (*pRbcS, pUM140_0016* or *pUM056_0044*) at 154 or 178bp before the TSS, respectively. As a positive control we used the *BleR* coding sequence with *RbcsI* incorporated as described in (Oertel et al., 2015), controlled by either *pRbcS, pUM140_0016* or *pUM056_0044*. Similar to the promoter screen, we transformed *Ulva* gametes with these constructs and counted the number of resistant individuals for every promoter-intron combination. We observed no, or a strongly reduced number of resistant individuals when *RbcsI* was incorporated in *pRbcS, pUM140_0016* or *pUM056_0044* (Fig. 2c). In contrast, combinations where the *RbcsI* is incorporated as a stand-alone sequence or as part of a linker sequence are functional and generate a similar number of resistant individuals compared to the positive controls (Fig. 2c).

### Two approaches for transgene expression in *Ulva*

An important feature for a molecular toolkit is to allow overexpression of a transgene. In the green algae *Chlamydomonas*, expressing an unselected gene-of-interest (GOI) is challenging, but different strategies have been developed to increase efficiency (Rasala et al., 2012; Onishi and Pringle, 2016). We tested two approaches for *Ulva*: expressing a GOI and a selectable marker (BleR) from two transcripts or as a single, bicistronic transcriptional unit by utilizing ribosome-skipping 2A peptides (Liu et al., 2017).

To express two transcripts from two promoters, we made expression vectors containing both the selection marker (*pRbcS-BleR-tRbcS* (Oertel et al., 2015)) and *pUM140_0016-GOI-NOST*. For each GOI (*mTagBFP2, YFP* or *mCherry*), a second expression vector was created that also included *RbcsI* upstream of TSS (Fig. 3; Fig. S3; Fig. S4). A second approach for transgene expression is using a bicistronic mRNA, where the BleR cassette is transcriptionally coupled to the transgene of interested via a sequence encoding a ribosome-skipping 2A peptide (Rasala et al., 2012; Liu et al., 2017). In such a vector composition, two or more genes are transcribed as one mRNA, but the 2A peptides mediate ribosome skipping during translation (Liu et al., 2017). To investigate this approach in *Ulva*, we fused the *BleR* cassette to *mTagBFP, YFP* or *mCherry* via four different 2A peptides: equine rhinitis A virus (E2A), foot and mouth disease virus 2A (F2A), porcine teschovirus-1 2A (P2A) or toshea asigna virus 2A (T2A; Fig. 3; Fig. S4; Fig. S5; (Liu et al., 2017)). In addition, we used two promoters (*pRbcS* and *pUM140_0016*) to control expression of the bicistronic transcript (Fig. 3; Fig. S4; Fig. S5). We transformed *Ulva* gametes with these vector combinations and selected transformants using phleomycin (Oertel et al., 2015).

**Figure 3.**
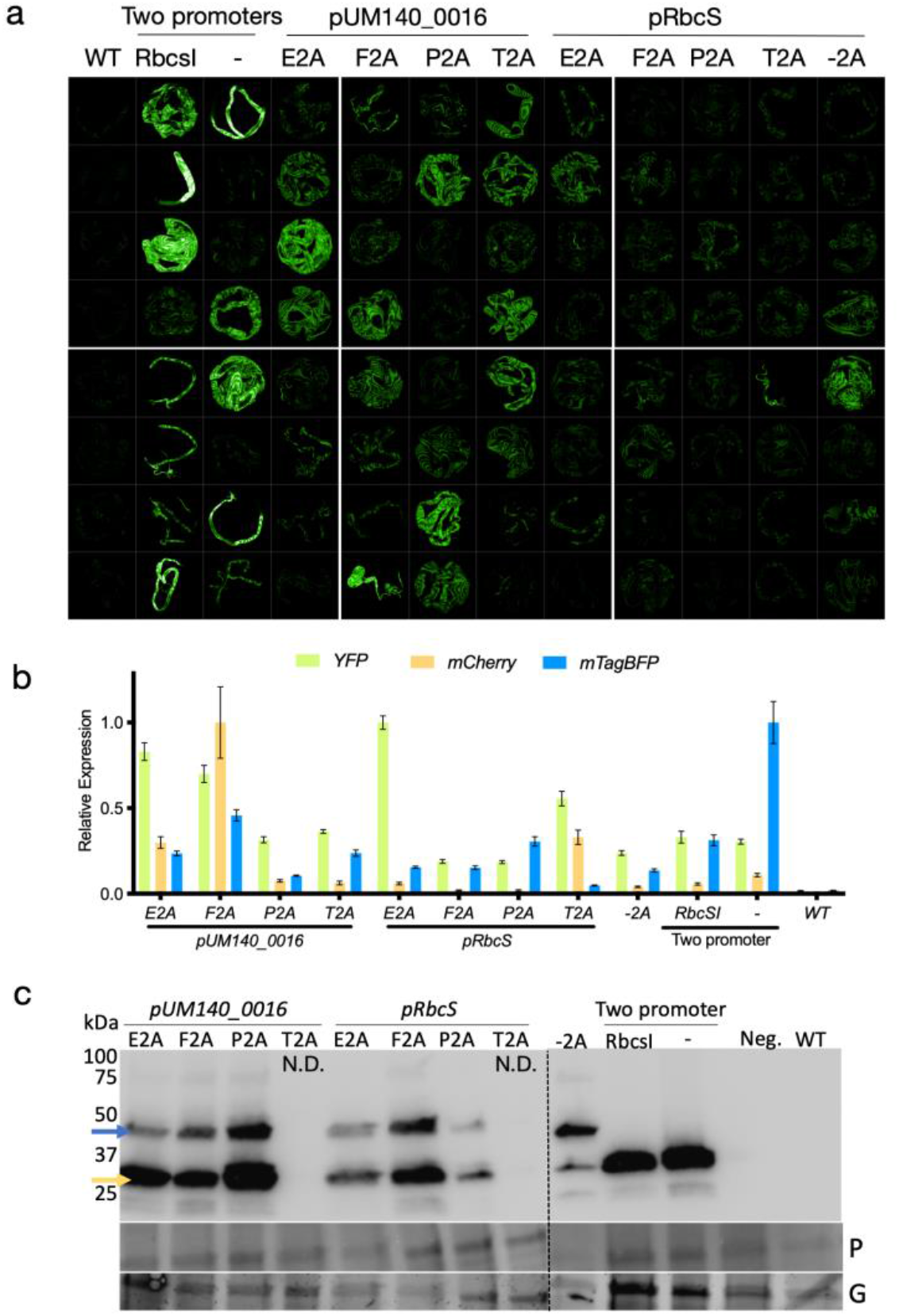
Stable expression of transgenes in *Ulva*. a) Confocal imaging visualizing the expression YFP in transgenic *Ulva* individuals and untransformed control (WT). *Ulva* gametes were transformed with expression vectors containing two promoters (*pRbcS-BleR-tRbcs* and *pUM140_0016-YFP-tNOS* with *RbcsI* or not (−)), one promoter (*pUM140_0016-BleR-2A-YFP-tRbcS* and *pRbcS-BleR-2A-YFP-tRbcS*) or a direct fusion to BleR (*pRbcS-BleR-YFP-tRbcS*; −2A). Eight primary transformants per construct are shown. See Figure S4 for constructs containing mCherry or mTagBFP2. b) qRT-PCR analysis of relative *YFP, mCherry, mTagBFP* expression in *Ulva* transgenic lines transformed with expression vectors containing two promoters (*pRbcS-BleR-tRbcs* and *pUM140 0016-GOI-tNOS*) or one promoter (*pRbcS-BleR-2A-GOI-tRbcS*) or a direct fusion to BleR (*pRbcS-BleR-GOI-tRbcS*; −2A) and untransformed control (WT). Values are mean of three technical replicates ± SE. c) Immunoblot analysis on *Ulva* transgenic lines transformed with expression vectors containing two promoters (*pRbcS-BleR-tRbcs* and *pUM140_0016-mCherry-tNOS*) or one promoter (*pRbcS-BleR-2A-mCherry-tRbcS*), a direct fusion of BleR-mCherry (−2A), vector control (Neg.) and untransformed control (WT). The expected size of the fusion protein (BleR-mCherry) and the cleaved mCherry is indicated with blue and yellow arrow, respectively. Loading control for each line is represented with ponceau staining (P) and stain-free gel imaging (G). N.D.: Not determined. See Figure S4 for constructs containing YFP or mTagBFP2.

Expression of transgenes was confirmed for all constructs by confocal microscopy, qRT-PCR and immunoblot analyses (Fig. 3; Fig. S3; Fig. S4; Fig. S5). For individuals expressing *YFP* using a two-promoter approach, we visualized expression in 67% and 66% of transformants in the presence or absence of *RbcsI* before the TSS, respectively (Fig. S3). Although there is variability in transgene expression between individuals, we see high and constitutive expression in ~25% of individuals. YFP signal was lower or confined to a subset of cells in other individuals (Fig. S3). Furthermore, we did not observe major differences in transgene expression when the vectors were delivered as linear or circular molecules (Fig. S3). The immunoblot analysis demonstrated that inclusion of E2A, F2A and P2A and T2A resulted in efficient separation of the GOI from BleR, although the fusion protein was detected as well for all lines (Fig. 3c; Fig. S4). Although no clear differences in RNA or protein accumulation are detected using qRT-PCR and immunoblot where pooled individuals were investigated; a bicistronic expression cassette under control of *pRbcS* appears to result in lower accumulation of fluorescent proteins compare to *pUM140_0016* (Fig. 3a; Fig. S4; Fig. S5). The transgene is stably integrated in the *Ulva* genome and inherited as expression was observed in the following generation (T1; Fig. S5). Building on these results, we made expression vectors containing BleR fused to F2A and one of three other genes encoding fluorescent proteins (*mCerulean, GFP* and *tdTomato* (Rasala et al., 2013)). We demonstrated overexpression of the transgenes using confocal imaging (Fig. S5). In conclusion, high overexpression of transgenes can be achieved in *Ulva* with one or two transcriptional units.

### Creating marker lines using transit peptides and transgenes

To facilitate the assembly of expression vectors containing endogenous or heterologous genes, we designed standard destination vectors where only the GOI and tag need to be inserted (Fig. S6; Materials and Methods). When two expression cassettes are preferred, we designed *pGG-pUM140_0016-BE-NOST-PIBT3* and *pGG-pUM140_0016-BE-NOST-PIBGT*, PIBGT and PIBT3 denote BleR fused to GFP, or not, and BE the Golden Gate overhang sites ((Lampropoulos et al., 2013); Fig. S6). Similarly, we designed pGG-*pUM140_0016-BleR-2A-CE-UmtRbcS* and pGG-*UmpRbcS-BleR-2A-CE-UmtRbcS* with the four 2A sequences (E2A, F2A, P2A and T2A). Additionally, we created destination vectors that allow straightforward assembly of more complex or multicistronic constructs (pGG-*pUM140_0016-BleR-2A-BG-UmtRbcS* and pGG-*UmpRbcS-BleR-2A-BG-UmtRbcS*) together with the necessary 2A entry vectors (Table S1; Fig. S6). The GOI with an N- and/or C-terminal tag will replace the *ccdB/*CmR counterselection module.

In summary, our molecular toolkit allows one to efficiently generate a diverse set of vectors for *Ulva* stable transformation. In total, 15 out of 20 *promoter-BleR-GFP-tRbcS* combinations were functional in *Ulva* and we identified two promoters that consistently result in more resistant individuals compared to the previously described *pRbcS*. A promoter deletion study indicated that the main transcriptional enhancers are located in the first 250bp of *pRbcS, pUM140_0016* and *pUM056 0044*. Furthermore, the importance of incorporating endogenous intron sequences in transgenes is highlighted by our intron-variant experiment. Using two expression systems, we can efficiently generate transgenic *Ulva* lines.

With the development of our molecular toolkit in hand, we tested the possibility of generating tagged *Ulva* lines. We selected transit peptides (TPs) functionally characterized in *Chlamydomonas* and endogenous *Ulva* genes for overexpression, as a bicistronic construct fused to BleR-F2A and as a second expression cassette, respectively. Thirteen TPs were cloned: six C-terminal peptides including three different nuclear localization signals (SV40, 2xSV40 and N7) and three markers for microbodies (Pumpkin Malate Synthase (PMS), Chlamydomonas Malate Synthase (CrMS) and Random Peroxisome Targeting Sequence (RMS)), five N-terminal tags including two peptides for mitochondrial targeting (HSP70C and atpA), two chloroplast targeting signals (psaD and USPA), one signal for excretion to the extracellular matrix (cCA) and two N-terminal endoplasmic reticulum (ER) targeting signals combined with a C-terminal HDEL ER-retention signal (ars1 and BIP; Table S1; Table S2; (Lampropoulos et al., 2013; Lauersen et al., 2015; Crozet et al., 2018)). We cloned the TPs in the appropriate entry clones and assembled them with YFP in pGG-*pUM140_0016-BleR-F2A-BG-UmtRbcS-KmR*. We observed *YFP* expression in 75-100% selected transformants for eleven out of thirteen constructs (Fig. 4a; Fig. S7). All constructs where no YFP signal was observed (atpA, BIP and USPA) still contain their endogenous *Chlamydomonas* intron, possibly suggesting that correct splicing is impaired in the transgene. HSP70c is the only transit peptide containing the *Chlamydomonas* intron that is functional. For atpA we observed YFP signal with anticipated localization when using the coding sequence (CDS; Fig. 4a; Fig. S7). For two TPs, cCA and psaD, we only observed free YFP signal, indicating that targeting to the extracellular matrix or chloroplast was impaired (Fig. 4a; Fig. S7). In contrast, we observed YFP expression consistent with localization in mitochondria (atpA (CDS) and HSP70c), microbodies (CMS, PMS and RMS), the nucleus (SV40, CrNLS and N7) and ER (ars1). Importantly, we observed no obvious nuclear localization as visualized in direct fusion of BleR with a fluorescent protein (Fig. S2b; Fig. S5; (Oertel et al., 2015)), suggesting that ribosome-skipping is efficient and no substantial amount of fusion protein is generated in these lines.

**Figure 4.**
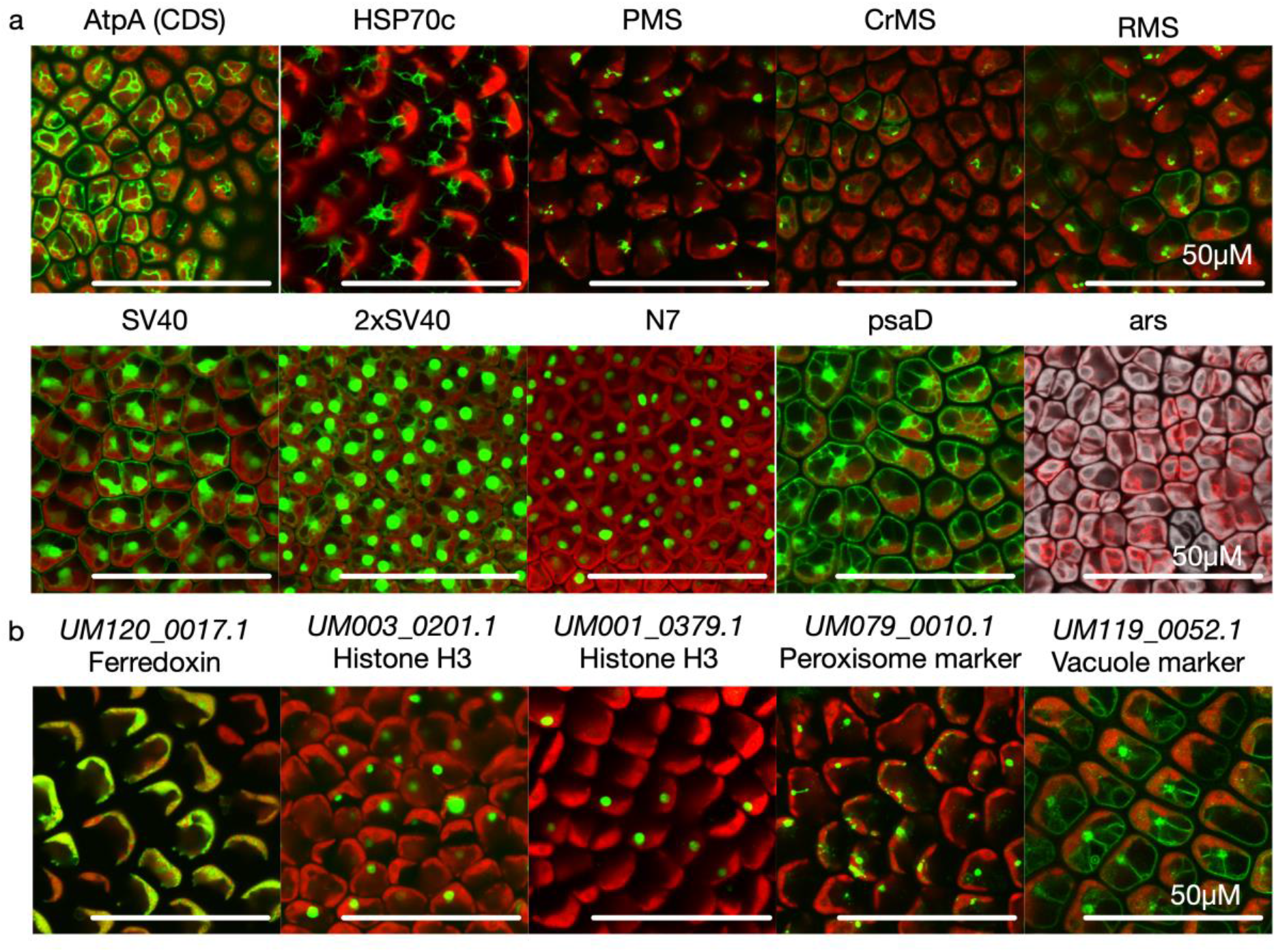
Tagged *Ulva* transgenic lines. a) Visualisation of *Ulva* transgenic lines expressing *YFP* or *mCherry* targeted to different intracellular locations using transit peptides: mitochondria (AtpA and HSP70c), microbodies (PMS, CrMS and RMS), nucleus (SV40; 2xSV40 and N7) and ER (ars-mCherryHDEL). One transgenic line (psaD) shows free YFP signal. For each line a representative individual is shown. Scale bar: 50μM. See also Figure S7. b) Visualisation of *Ulva* transgenic lines expressing endogenous *Ulva* genes tagged with YFP targeted to different intracellular locations: chloroplast (UM120_0017.1), nucleus (UM003_0201.1 and UM001_0379.1), microbodies (UM079_0010.1) and vacuole (UM110_0052.1). For each line a representative individual is shown. Scale bar: 50μM. See also Figure S8.

Besides small transit peptides, we identified twenty *Ulva* orthologs of *Chlamydomonas* marker genes for different intracellular locations (nucleus, mitochondria, chloroplast, ER, peroxisomes; (Mackinder et al., 2017)). We cloned eleven genes from genomic DNA in the appropriate entry vector and cured BsaI sites where necessary (Table S3). Genomic sequences containing endogenous introns were chosen based on our and other data that these introns could facilitate transgene expression (Fig. 2c, (Lumbreras et al., 1998)). We assembled the gene sequences in destination vector *pGG-pUM140_0016-BE-NOST-PIBT3-* KmR and included a C-terminal YFP. *Ulva* gametes were transformed and *YFP* expression was visualized in blade cells. We observed expected nuclear localization for two tagged histone H3 proteins (encoded by *UM001_0379* and *UM003_0201*), peroxisomal localization for the tagged protein encoded by an ortholog (*UM079_0010*) of the peroxisomal marker encoded by the *Chlamydomonas* gene *CR10G07890*, UM119_0052-YFP was localized in vacuole membranes and the tagged Ferredoxin ortholog (encoded by *UM120_0017*) localizes in *Ulva* chloroplasts (Fig. 4b; Fig. S8). Furthermore, we were able to generate tagged lines for a *Ulva* orthologs coding for α-tubulin (*TUA2; UM021_0090’*), EF-hand-containing Ca2+/calmodulin-dependent protein kinase (EF; *UM005_0221*), 20 kDa translocon at the inner membrane of chloroplasts (TIC20; *UM014_0107*) and Translocase of chloroplast 34 (TOC34; UM025_*0020*; Fig. S8). No or unclear localization was observed for the remaining two tagged genes.

In conclusion, we illustrate that a variety of *Ulva* marker lines can be generated by fusing transit peptides or endogenous proteins to a fluorescent marker.

## Discussion

*U. mutabilis* is a model organism for multicellular green seaweeds. With the completion of the *Ulva* genome project (De Clerck et al., 2018), the development of a flexible molecular toolkit is required for continued functional genomic research in this species. Prior to this report, stable transgene expression in *Ulva* had only been demonstrated for the BleR cassette, allowing selection of transformants (Oertel et al., 2015). The original vectors contain multiple cloning sites that permit the creation of larger expression vectors containing more modules (Oertel et al., 2015). However, to allow more uniform and modular assembly of expression vectors, we opted to base our toolkit on the GreenGate cloning system (Lampropoulos et al., 2013). Our toolkit differs from other MoClo-based modular kits generated for *Chlamydomonas* and Cyanobacteria (Crozet et al., 2018; Vasudevan et al., 2019) that allow fusion of eleven sites. The GreenGate strategy contains six modules (Lampropoulos et al., 2013), which implies a more straightforward design for assembly of common expression vectors, but still permits to generate more complex constructs such as gene stacking (Lampropoulos et al., 2013). In this report, we demonstrate the efficient generation of transgenic *Ulva* lines expressing transgenes, a feature that has not been achieved before in other multicellular algae. *Ulva* therefore has the potential to complement unicellular model algae such as *Chlamydomonas* as a workhorse for functional genetic research and as a chassis for synthetic biology.

Although it remains unclear what sequence characteristics are indispensable for a functional promoter in (green) algae, we report 15 functional constitutive promoters for *Ulva* with apparent varying degrees of activity. Interestingly, promoter regions of homologous genes between *Ulva* and *Chlamydomonas* are functional. Not only the endogenous promoter of the small subunit of rubisco gene are functional both in *Ulva* and *Chlamydomonas*, but also promoter regions from *PSAD* and two genes encoding chlorophyll a/b binding proteins (Blankenship and Kindle, 1992; Fischer and Rochaix, 2001). Additional promoter screens to identify conditional promoters that are only active upon addition of a stimulus (*e.g*. light or heat shock), in specific cell types or during specific developmental phases would be a useful addition to the collection we report here.

Our study confirms the importance of inserting an endogenous intron sequence in the BleR resistance cassette (Lumbreras et al., 1998; Oertel et al., 2015). Importantly, *RbcsI* does not necessarily need to be introduced in BleR to promote transgene expression but can be placed immediately before the TSS. This implies that transcriptional enhancers do not necessarily need to be built into a coding sequence to increase transgene expression. However, in contrast to *Chlamydomonas* (Lumbreras et al., 1998), we did not observe an increase in transformation efficiency when *RbscI* was included more upstream in a promoter sequence. The endogenous promoter *pUM140_0016* promotes transgene expression in a vector system containing two expression cassettes without the addition of *RbscI*, suggesting that all necessary regulatory and enhancer elements are present.

We developed two vector systems for transgene expression in *Ulva*, one containing two expression cassettes and one for bi- or multicistronic cassettes. Our vector containing two expression cassettes generates more transgenic individuals with detectable transgene expression compared to the model *Chlamydomonas* (67% versus 13% (Onishi and Pringle, 2016)). In a bicistronic context, the transgene is forced to be transcribed together with BleR. For the creation of bicistronic expression cassettes, we tested four different 2A ribosome-skipping peptides but alternative approaches exist, such as the addition of internal ribosome entry site (IRES) or even short linkers of unstructured sequences between the selectable marker and the GOI (Onishi and Pringle, 2016). Here, we chose to characterize 2A peptides as this was reported to be compatible with BleR expression in *Chlamydomonas*, whereas IRES or short linkers were not (Rasala et al., 2012; Onishi and Pringle, 2016). Although both vector systems allow high expression of transgenes in *Ulva*, immunoblot analysis suggests that while ribosome cleavage is efficient, it is not perfect. As a consequence, we advise to first attempt overexpression using two expression cassettes.

We successfully developed 18 *Ulva* marker lines for six intracellular locations (chloroplast, endoplasmic reticulum, microbodies, mitochondria, nucleus and vacuole). These are important references to investigate the intracellular localization of tagged genes. We generated multiple independent lines targeting three intracellular compartments. Our results suggest that N7, AtpA (CDS) and PMS result in highly specific localization in nucleus, mitochondria and microbodies, respectively, whereas more aspecific YFP signal was observed in transgenic lines expressing other TPs. Importantly, we demonstrate that eight transit peptides tested are functional both in *Chlamydomonas* and *Ulva*. Furthermore, we illustrate *Ulva* genes orthologous to known *Chlamydomonas* marker genes show intracellular localization consistent with their proposed function, *e.g*. nuclear localization for two proteins encoding a histone H3 protein. With the generation of these lines, we report the first tagged endogenous genes described in *Ulva*. Generating transgenic lines with tagged transgenes allows the inspection of intracellular localization, but also enables the development of techniques such as Chromatin Immunoprecipitation or affinity purification-mass spectrometry.

We are currently able to generate gain-of-function mutants in *Ulva*. Although the generation of T-DNA insertion mutants has been described (Oertel et al., 2015), the development of RNAi techniques or programmable genome editing techniques such as CRISPR/Cas9 would be an important addition to the molecular toolkit of *Ulva*. The overall low efficiency of CRISPR/Cas9 reported in algae and the single functional selection marker (BleR) in *Ulva* make the development of this technique challenging at this time.

In summary, the toolkit described here allows the generation of *Ulva* transgenic lines stably overexpressing transgenes. We illustrate this with the expression of fluorescent proteins, marker lines and tagged endogenous proteins. To our knowledge, *Ulva* is the first seaweed that is amendable for these experiments at this time. Besides enabling molecular studies in relevant processes for *Ulva* biology (De Clerck et al., 2018), the homologs of genes from model microalgae such as *Chlamydomonas* can now be studied in a multicellular context.

## Materials and Methods

### Algae growth and cultivation

*Ulva mutabilis* Føyn “slender” mutant strain is a direct descendant of the original isolates of B. Føyn from the South Atlantic coast of Portugal near Ohlão and Faro (Føyn, 1958). The untransformed individuals were kindly provided by Thomas Wichard (University of Jena, Germany). Individuals were cultivated in tissue culture chamber under long day conditions (16h light:8h dark; 21°C; 75μM; Spectralux Plus NL-T8 36W/840/G13 fluorescent lamp) in standard Petri dishes containing synthetic Ulva Culture Medium (UCM; (Stratmann et al., 1996; Boesger et al., 2018)). *Ulva* was grown and parthenogenetically propagated as described previously (Wichard and Oertel, 2010).

### *Ulva* transformation

*Ulva* gametes were transformed using the PEG-mediated protocol described in (Oertel et al., 2015; Boesger et al., 2018). For each transformation, 3-5μg of plasmid DNA was used per construct. Plasmids were purified using a GeneJet Plasmid Miniprep kit (Thermo Scientific). 50μg/mL Phleomycin (Invivogen) was added 48h after transformation, resistant germlings develop parthenogenetically from the transformed gametes and can be detected after 2-3 weeks of cultivation. For each transformation experiment, non-vector controls were included to confirm proper selection. For experiments where relative transformation efficiency was scored, all constructs and controls were transformed in the same transformation experiment to reduce variability in transformation efficiency between different batches of gametes and to ensure the same input DNA was delivered in the same number of input gametes. Resistant individuals were counted and removed from the transformation plates; additional selection medium was added to allow detection of slower growing germlings. Mutant cultures are preserved and are available upon request.

### Vector assembly

Vector assembly is based on the GreenGate cloning system (Lampropoulos et al., 2013). Sequences for entry modules were obtained via gene synthesis (BioXP3200, Codex DNA) or PCR-amplification of the target DNA sequence using primers that contain a ~20bp overlap with the entry module using high-fidelity polymerase Q5 (NEB) or GXL (Takara Bio). After gel electrophoresis, the amplicon is purified (Zymoclean Gel DNA Recovery Kit, Zymo Research) and mixed with the pre-digested (BsaI, Thermo Scientific) entry clone and NEBuilder (NEB). After assembly (15-60 minutes at 50°C), the mix is transformed into chemically-competent DH5α *Escherichia coli* cells and plated out on LB+100μg/mL Carbenicillin. PCR-mediated silent mutations were introduced to remove internal BsaI recognition sites. All template plasmids containing genes encoding fluorescent proteins or transit peptides were ordered via Chlamydomonas Research Center (https://www.chlamycollection.org; Table S1, Table S2). All *Ulva* gene and promoter sequences were cloned from genomic DNA, isolated via CTAB (Clarke, 2009) method or Omniprep for plant (G-Biosciences).

For a Golden Gate reaction, 100ng of each entry and destination vector are assembled in one reaction mix containing 10U BsaI-HF-v2 (NEB), 200U T4 Ligase (NEB), 1mM ATP (Thermo Scientific) and 1x Cutsmart buffer (NEB). The reaction mix is incubated at 30x(18°C (3 minutes) and 37°C (3 minutes)), followed by 5 minutes 50°C and 5 minutes 80°C. After assembly, the mix is transformed in DH5α *E. coli* cells and plated out on LB+25μg/mL Kanamycin. Vectors were validated using Sanger sequencing (Mix2seq, Eurofins) and restriction digest, typically using NcoI (Promega), PstI (Promega) or XhoI (NEB).

All entry clones in this study are designed as described in (Lampropoulos et al., 2013). The destination vector used is a high-copy number pGG-AG-KmR. The ccdB/CmR cassette from pEN-L1-AG-L2 (Houbaert et al., 2018) was amplified using pGG-AG-KmR primers (Table S4). The PCR product was purified and DNA was digested by ApaI (Promega), then the digest was purified. The pEN-L1-AG-L2 vector was digested by EcoRV and ApaI (Promega) and purified from gel. The vector fragment was ligated to the ApaI digested PCR product to generate pGG-AG-KmR. For vectors with two expression cassettes, the BleR resistance cassette was inserted proximal to the G BsaI recognition site. The pGG-AG-KmR was digested using EcoRV and NotI (Promega) and purified from gel, the PIBT3 or PIBGT cassettes were amplified from pPIBT3 or pPIBGT8 (Oertel et al., 2015) using pGG-AG-PIBT-KmR primers (Table S4) and gel purified. The amplicons were inserted in the digested pGG-AG-KmR vector via Gibson Assembly to generate pGG-AG-PIBT3-KmR or pGG-AG-PIBGT-KmR, where PIBGT contains a BleR-GFP fusion.

For the construction of modified destination vectors, first a Golden Gate reaction was performed containing all desired fragments and two or three variable linkers containing AarI sites and the SacB counterselectable marker (Decaestecker et al., 2019). After sequence verification, the vectors were digested with AarI and purified from gel. The ccdB/CmR cassette from pEN-L1-AG-L2 (Houbaert et al., 2018) was amplified using B-ccdB/CmR-G or C-ccdB/CmR-E primers (Table S4). The amplicons were inserted in the digested vectors vector via Gibson Assembly to generate modified destination vectors for two expression cassettes (pGG-pUM140_0016-BE-NOST-PIBT3 or pGG-pUM140_0016-BE-NOST-PIBGT) or one expression cassette (pRbcS-BleR-2A-CE-tRbcS, pRbcS-BleR-2A-BG-tRbcS, pUM140_0016-BleR-2A-CE-tRbcS or pUM140_0016-BleR-2A-BG-tRbcS containing E2A, F2A, P2A or T2A; Fig. S7). All destination vectors are transformed in One Shot ccdB Survival 2 T1R Chemically Competent Cells (ThermoFisher).

All vectors generated in this study (Table S1) are available at https://gatewayvectors.vib.be.

### Confocal imaging

Individual thalli of one-month-old resistant individuals were selected for imaging. Entire individuals were imaged using an Opera Phenix High-Content Screening System (Perkin Elmer) and analyzed using Columbus software. Detailed imaging of tissues and cells was performed using Olympus Fluoview (FV1000) confocal microscope with Fluoview software. The following excitation and emission settings were used: YFP, 515 excitation with 530-545 emission; mCherry, 559 excitation with 564-644 emission; mTagBFP2 405 excitation with 446-546 emission and chlorophyll, 559 excitation with 650-750 emission. For each transgenic line, four to ten independent individuals were screened, a representative image is selected for all lines with a functional construct.

### qRT-PCR

One-month-old resistant individuals were harvested, flash frozen and RNA was extracted using the ReliaPrep^™^ RNA Tissue Miniprep System (Promega) according to the manufacturer’s specifications. qRT-PCR experiments were performed in a Light-Cycler480 Real-Time SYBRgreen PCR system (Roche). 500 ng of RNA was reverse-transcribed with the qScript^®^ cDNA Supermix kit (QuantaBio) according to the manufacturer’s specifications. qRT-PCR results were normalized against two reference genes (*EFLa (UM025_0053*) and *UBQ10* (*UM008_0183*)). For each experiment, three technical replicates were performed on the same biological replicate. Each biological replicate consists of multiple individuals grown under the same conditions; harvesting and further sample handling were done in separate tubes for the indicated transgenic lines. All primers used in this study are listed in Table S4.

### Protein extraction and Immuno Blot

Protein extracts were extracted from different transgenic lines using in a 2/3 ratio (v/w) of extraction buffer (25 mm Tris-HCl, pH 7.6, 15 mm MgCl2, 150 mm NaCl, 15 mm p-nitrophenyl phosphate, 60 mm β-glycerophosphate, 0.1% (v/v) Nonidet P-40, 0.1 mm sodium vanadate, 1 mm NaF, 1 mm PMSF, 1 μm trans-epoxysuccinyl-l-leucylamido-(4-guanidino)butane (E64), 5% (v/v) ethylene glycol; EDTA-free Ultra Complete Tablet (Sigma)). After two freeze-thaw cycles in extraction buffer, the soluble protein fraction was obtained by a two centrifugation steps at 14000rpm for 15 minutes 4 °C. Concentration was determined via Bradford assay (Bio-rad). 15-20μg protein was loaded on a 4-15% Mini-PROTEAN TGX Precast gel (Bio-Rad). After SDS-PAGE, proteins were transferred to PVDF membrane using Trans-Blot Turbo Transfer System (Bio-Rad). Correct transfer was visualized using Ponceau Staining. After destaining, an excess of 3% SM-milk in TBT was applied to block the membrane (RT, 20 min). The blot was incubated overnight on a shaker (4°C) with the primary antibody (anti-GFP 3H9, anti-RFP 6G6 (Chromotek) or anti-tRFP (Evrogen)). After three washes with TBT buffer, the secondary HRP-conjugated antibody was added (anti-mouse A9917 (Sigma) or anti-rabbit GENA934 (Sigma); RT, 1h). Detection was performed using Clarity Western ECL Substrate (Bio-Rad) and a ChemiDoc MP System (Bio-Rad). Each biological replicate consists of multiple individuals grown under the same conditions; harvesting and further sample handling were done in separate tubes for the indicated transgenic lines.

## Large datasets

All generated vectors of the toolkit (Table S1) are available on https://gatewayvectors.vib.be. Transgenic lines are maintained and are available from the corresponding author upon reasonable request.

## Acknowledgements

J.B. thanks the Research Foundation – Flanders (FWO) and Ghent University for a postdoctoral fellowship (12T3418N and BOF20/PDO/016). X.L. is indebted to the China Scholarship Council (201504910698) and Ghent University Special Research Fund (BOF-16/CHN/023) for a PhD grant. The authors thank Thomas Wichard and Michiel Kwantes (University of Jena) for providing pPIBT3 and pPIBGT8 vectors and untransformed *Ulva mutabilis* individuals. The VIB Screening Core facility assisted in imaging entire transgenic individuals and image analysis. Mansour Karimi (VIB-University of Ghent) designed and created the pGG-AG-KmR destination vector.

## Author contribution

J.B., T.J. and O.D.C. designed the study. J.B. performed all experiments and analyzed the data. X.L. maintained *Ulva* cultures for experiments. J.B. wrote the manuscript with contributions from all authors. O.D.C agrees to serve as the author responsible for contact and ensures communication.

## Supporting Information

**Figure S1.** Promoter screen gene identifiers and cloning strategy.

**Figure S2.** Promoter screen expression analysis

**Figure S3.** Visualization of YFP expression in *Ulva* transgenic lines expressing *pUM140_0016-RbcsI-YFP-NOST* or *pUM140_0016-YFP-NOST*.

**Figure S4.** Expression analysis of transgenic *Ulva* lines.

**Figure S5.** Expression analysis of transgenic *Ulva* lines using bicistronic expression vectors.

**Figure S6.** Custom destination vectors cloning scheme.

**Figure S7.** Expression analysis of transgenic *Ulva* marker lines using transit peptides.

**Figure S8.** Expression analysis of transgenic *Ulva* marker lines using endogenous genes.

**Table S1.** Complete list of generated vectors (entry, destination and expression).

**Table S2.** Selection of transit peptides.

**Table S3.** Selection of *Ulva* marker genes.

**Table S4.** List of primers used in this study.

## Parsed Citations

**Blaby-Haas CE, Merchant SS (2019) Comparative and Functional Algal Genomics. Annual Review of Plant Biology 70: 605–638**

Google Scholar: <u>Author Only Title Only Author and Title</u>

**Blankenship JE, Kindle KL (1992) Expression of chimeric genes by the light-regulated cabII-1 promoter in Chlamydomonas reinhardtii: a cabII-1/nit1 gene functions as a dominant selectable marker in a nit1-nit2-strain. Molecular and Cellular Biology 12: 5268–5279**

Google Scholar: <u>Author Only Title Only Author and Title</u>

**Boesger J, Kwantes M, Wichard T, Kwantes M, Wichard T (2018) Polyethylene glycol-mediated transformation in the green macroalga Ulva mutabilis (Chlorophyta): A forward genetics approach. Protocols for Macroalgae Research 469–483**

Google Scholar: <u>Author Only Title Only Author and Title</u>

**Bolton JJ, Cyrus MD, Brand MJ, Joubert M, Macey BM (2016) Why grow Ulva? Its potential role in the future of aquaculture.**

**Perspectives in Phycology 113–120**

Google Scholar: <u>Author Only Title Only Author and Title</u>

**Cai C, Wang L, Jiang T, Zhou L, He P, Jiao B (2017) The complete mitochondrial genomes of green tide algae Ulva flexuosa (Ulvophyceae, Chlorophyta). Conservation Genet Resour 1–4**

Google Scholar: <u>Author Only Title Only Author and Title</u>

**Casini, A Storch M, Baldwin GS, Ellis T (2015) Bricks and blueprints: methods and standards for DNA assembly. Nature Reviews Molecular Cell Biology 16: 568–576**

Google Scholar: <u>Author Only Title Only Author and Title</u>

**Charrier B, Abreu MH, Araujo R, Bruhn Ą Coates JC, De Clerck O, Katsaros C, Robaina RR, Wichard T (2017) Furthering knowledge of seaweed growth and development to facilitate sustainable aquaculture. New Phytol 216: 967–975**

Google Scholar: <u>Author Only Title Only Author and Title</u>

**Clarke JD (2009) Cetyltrimethyl Ammonium Bromide (CTAB) DNA Miniprep for Plant DNA Isolation. Cold Spring Harb Protoc 2009: pdb.prot5177**

Google Scholar: <u>Author Only Title Only Author and Title</u>

**Cock JM, Sterck L, Rouzé P, Scornet D, Allen AE, Amoutzias G, Anthouard V, Artiguenave F, Aury J-M, Badger JH, et al (2010) The Ectocarpus genome and the independent evolution of multicellularity in brown algae. Nature 465: 617–621**

Google Scholar: <u>Author Only Title Only Author and Title</u>

**Collén J, Porcel B, Carré W, Ball SG, Chaparro C, Tonon T, Barbeyron T, Michel G, Noel B, Valentin K, et al (2013) Genome structure and metabolic features in the red seaweed Chondrus crispus shed light on evolution of the Archaeplastida. PNAS 110: 5247–5252**

Google Scholar: <u>Author Only Title Only Author and Title</u>

**Crozet P, Navarro FJ, Willmund F, Mehrshahi P, Bakowski K, Lauersen KJ, Pérez-Pérez M-E, Auroy P, Gorchs Rovira A Sauret-Gueto S, et al (2018) Birth of a Photosynthetic Chassis: A MoClo Toolkit Enabling Synthetic Biology in the Microalga Chlamydomonas reinhardtii. ACS Synthetic Biology 7: 2074–2086**

Google Scholar: <u>Author Only Title Only Author and Title</u>

**De Clerck O, Kao S-M, Bogaert KA, Blomme J, Foflonker F, Kwantes M, Vancaester E, Vanderstraeten L, Aydogdu E, Boesger J, et al (2018) Insights into the Evolution of Multicellularity from the Sea Lettuce Genome. Current Biology 28: 2921–2933.e5**

Google Scholar: <u>Author Only Title Only Author and Title</u>

**Decaestecker W, Buono RA, Pfeiffer ML, Vangheluwe N, Jourquin J, Karimi M, Isterdael GV, Beeckman T, Nowack MK, Jacobs TB (2019) CRISPR-TSKO: A Technique for Efficient Mutagenesis in Specific Cell Types, Tissues, or Organs in Arabidopsis. The Plant Cell 31: 2868–2887**

Google Scholar: <u>Author Only Title Only Author and Title</u>

**Doron L, Segal N, Shapira M (2016) Transgene Expression in Microalgae-From Tools to Applications. Front Plant Sci. doi: 10.3389/fpls.2016.00505**

Google Scholar: <u>Author Only Title Only Author and Title</u>

**Faktorová D, Nisbet RER, Robledo JAF, Casacuberta E, Sudek L, Allen AE, Ares M, Aresté C, Balestreri C, Barbrook AC, et al (2020) Genetic tool development in marine protists: emerging model organisms for experimental cell biology. Nat Methods 17: 481–494**

Google Scholar: <u>Author Only Title Only Author and Title</u>

**Fischer N, Rochaix J-D (2001) The flanking regions of PsaD drive efficient gene expression in the nucleus of the green alga Chlamydomonas reinhardtii. Mol Gen Genomics 265: 888–894**

Google Scholar: <u>Author Only Title Only Author and Title</u>

**Føyn B (1958) Über die Sexualität und den Generationswechsel von Ulva mutabilis. Arch Protistenkd 102: 473–480**

Google Scholar: <u>Author Only Title Only Author and Title</u>

**Fuhrmann M, Oertel W, Hegemann P (1999) A synthetic gene coding for the green fluorescent protein (GFP) is a versatile reporter in Chlamydomonas reinhardtii†. The Plant Journal 19: 353–361**

Google Scholar: <u>Author Only Title Only Author and Title</u>

**Houbaert A Zhang C, Tiwari M, Wang K, de Marcos Serrano *A*, Savatin DV, Urs MJ, Zhiponova MK, Gudesblat GE, Vanhoutte I, et al (2018) POLAR-guided signalling complex assembly and localization drive asymmetric cell division. Nature 563: 574–578**

Google Scholar: <u>Author Only Title Only Author and Title</u>

**Huang X, Weber JC, Hinson TK, Mathieson AC, Minocha SC (1996) Transient Expression of the GUS Reporter Gene in the Protoplasts and Partially Digested Cells of Ulva lactuca L. (Chlorophyta). Botanica Marina 39: 467–474**

Google Scholar: <u>Author Only Title Only Author and Title</u>

**Ichihara K, Yamazaki T, Miyamura S, Hiraoka M, Kawano S (2019) Asexual thalli originated from sporophytic thalli via apomeiosis in the green seaweed Ulva. Sci Rep 9: 1–12**

Google Scholar: <u>Author Only Title Only Author and Title</u>

**Jaeger D, Baier T, Lauersen KJ (2019) Intronserter, an advanced online tool for design of intron containing transgenes. Algal Research 42: 101588**

Google Scholar: <u>Author Only Title Only Author and Title</u>

**Jiang P, Qin S, Tseng CK (2003) Expression of the lacZ reporter gene in sporophytes of the seaweed Laminaria japonica (Phaeophyceae) by gametophyte-targeted transformation. Plant Cell Rep 21: 1211–1216**

Google Scholar: <u>Author Only Title Only Author and Title</u>

**Kakinuma M, Ikeda M, Coury DA, Tominaga H, Kobayashi I, Amano H (2009) Isolation and characterization of the rbcS genes from a sterile mutant of Ulva pertusa (Ulvales, Chlorophyta) and transient gene expression using the rbcS gene promoter. Fish Sci 75: 1015– 1028**

Google Scholar: <u>Author Only Title Only Author and Title</u>

**Lampropoulos A, Sutikovic Z, Wenzl C, Maegele I, Lohmann JU, Forner J (2013) GreenGate - A Novel, Versatile, and Efficient Cloning System for Plant Transgenesis. PLOS ONE 8: e83043**

Google Scholar: <u>Author Only Title Only Author and Title</u>

**Lauersen KJ, Kruse O, Mussgnug JH (2015) Targeted expression of nuclear transgenes in Chlamydomonas reinhardtii with a versatile, modular vector toolkit. Appl Microbiol Biotechnol 99: 3491–3503**

Google Scholar: <u>Author Only Title Only Author and Title</u>

**Lim J-M, Ahn J-W, Hwangbo K, Choi D-W, Park E-J, Hwang MS, Liu JR, Jeong W-J (2013) Development of cyan fluorescent protein (CFP) reporter system in green alga Chlamydomonas reinhardtii and macroalgae Pyropia sp. Plant Biotechnol Rep 7: 407–414**

Google Scholar: <u>Author Only Title Only Author and Title</u>

**Lim J-M, Shin YJ, Jung S, Choi SA, Choi D-W, Min SR, Jeong W-J (2019) Loss of copy number and expression of transgene during meiosis in Pyropia tenera. Plant Biotechnol Rep 13: 653–661**

Google Scholar: <u>Author Only Title Only Author and Title</u>

**Liu Z, Chen O, Wall JBJ, Zheng M, Zhou Y, Wang L, Vaseghi HR, Qian L, Liu J (2017) Systematic comparison of 2A peptides for cloning multi-genes in a polycistronic vector. Sci Rep 7: 1–9**

Google Scholar: <u>Author Only Title Only Author and Title</u>

**Loureiro R, Gachon CMM, Rebours C (2015) Seaweed cultivation: potential and challenges of crop domestication at an unprecedented pace. New Phytologist 206: 489–492**

Google Scholar: <u>Author Only Title Only Author and Title</u>

**Lumbreras V, Stevens DR, Purton S (1998) Efficient foreign gene expression in Chlamydomonas reinhardtii mediated by an endogenous intron. The Plant Journal 14: 441–447**

Google Scholar: <u>Author Only Title Only Author and Title</u>

**Mackinder LCM, Chen C, Leib RD, Patena W, Blum SR, Rodman M, Ramundo S, Adams CM, Jonikas MC (2017) A Spatial Interactome Reveals the Protein Organization of the Algal CO2-Concentrating Mechanism. Cell 171: 133-147.e14**

Google Scholar: <u>Author Only Title Only Author and Title</u>

**Melton III JT, Leliaert F, Tronholm A Lopez-Bautista JM (2015) The Complete Chloroplast and Mitochondrial Genomes of the Green Macroalga Ulva sp. UNA00071828 (Ulvophyceae, Chlorophyta). PLOS ONE 10: e0121020**

Google Scholar: <u>Author Only Title Only Author and Title</u>

**Norris SR, Meyer SE, Callis J (1993) The intron of Arabidopsis thaliana polyubiquitin genes is conserved in location and is a quantitative determinant of chimeric gene expression. Plant Mol Biol 21: 895–906**

Google Scholar: <u>Author Only Title Only Author and Title</u>

**Odell JT, Nagy F, Chua N-H (1985) Identification of DNA sequences required for activity of the cauliflower mosaic virus 35S promoter. Nature 313: 810–812**

Google Scholar: <u>Author Only Title Only Author and Title</u>

**Oertel W, Wichard T, Weissgerber A (2015) Transformation of Ulva mutabilis (Chlorophyta) by vector plasmids integrating into the genome. J Phycol 51: 963–979**

Google Scholar: <u>Author Only Title Only Author and Title</u>

**Onishi M, Pringle JR (2016) Robust Transgene Expression from Bicistronic mRNA in the Green Alga Chlamydomonas reinhardtii. G3 (Bethesda) 6: 4115–4125**

Google Scholar: <u>Author Only Title Only Author and Title</u>

**Rasala BA, Barrera DJ, Ng J, Plucinak TM, Rosenberg JN, Weeks DP, Oyler GA, Peterson TC, Haerizadeh F, Mayfield SP (2013) Expanding the spectral palette of fluorescent proteins for the green microalga Chlamydomonas reinhardtii. The Plant Journal 74: 545–556**

Google Scholar: <u>Author Only Title Only Author and Title</u>

**Rasala BA, Lee PA, Shen Z, Briggs SP, Mendez M, Mayfield SP (2012) Robust Expression and Secretion of Xylanase1 in Chlamydomonas reinhardtii by Fusion to a Selection Gene and Processing with the FMDV 2A Peptide. PLOS ONE 7: e43349**

Google Scholar: <u>Author Only Title Only Author and Title</u>

**Son SH, Ahn J-W, Uji T, Choi D-W, Park E-J, Hwang MS, Liu JR, Choi D, Mikami K, Jeong W-J (2012) Development of an expression system using the heat shock protein 70 promoter in the red macroalga, Porphyra tenera. J Appl Phycol 24: 79–87**

Google Scholar: <u>Author Only Title Only Author and Title</u>

**Song Q, Daozhan Y, Peng J, Changying T, Chengkui Z (2003) Stable Expression of lacZ Reporter Gene in Seaweed Undaria pinnatifida. Gaojishu Tongxun 13: 87–89**

Google Scholar: <u>Author Only Title Only Author and Title</u>

**Spoerner M, Wichard T, Bachhuber T, Stratmann J, Oertel W (2012) Growth and Thallus Morphogenesis of Ulva mutabilis (Chlorophyta) Depends on A Combination of Two Bacterial Species Excreting Regulatory Factors. J Phycol 48: 1433–1447**

Google Scholar: <u>Author Only Title Only Author and Title</u>

**Stratmann J, Paputsoglu G, Oertel W (1996) Differentiation of Ulva Mutabilis (chlorophyta) Gametangia and Gamete Release Are Controlled by Extracellular Inhibitors! Journal of Phycology 32: 1009–1021**

Google Scholar: <u>Author Only Title Only Author and Title</u>

**Strenkert D, Schmollinger S, Gallaher SD, Salomé PA, Purvine SO, Nicora CD, Mettler-Altmann T, Soubeyrand E, Weber APM, Lipton MS, et al (2019) Multiomics resolution of molecular events during a day in the life of Chlamydomonas. PNAS 116: 2374–2383**

Google Scholar: <u>Author Only Title Only Author and Title</u>

**Suzuki R, Yamazaki T, Toyoda A Kawano S (2014) A Transformation System Using rbcS N-Terminal Region Fused with GFP Demonstrates Pyrenoid Targeting of the Small Subunit of RubisCO in Ulva compressa. Cytologia 79: 427–428**

Google Scholar: <u>Author Only Title Only Author and Title</u>

**Uji T, Hirata R, Fukuda S, Mizuta H, Saga N (2014) A Codon-Optimized Bacterial Antibiotic Gene Used as Selection Marker for Stable Nuclear Transformation in the Marine Red Alga Pyropia yezoensis. Mar Biotechnol 16: 251–255**

Google Scholar: <u>Author Only Title Only Author and Title</u>

**Vasudevan R, Gale GAR, Schiavon AA, Puzorjov A, Malin J, Gillespie MD, Vavitsas K, Zulkower V, Wang B, Howe CJ, et al (2019) CyanoGate: A Modular Cloning Suite for Engineering Cyanobacteria Based on the Plant MoClo Syntax. Plant Physiology 180: 39–55**

Google Scholar: <u>Author Only Title Only Author and Title</u>

**Wang S, Li L, Li H, Sahu SK, Wang H, Xu Y, Xian W, Song B, Liang H, Cheng S, et al (2020) Genomes of early-diverging streptophyte algae shed light on plant terrestrialization. Nature Plants 6: 95–106**

Google Scholar: <u>Author Only Title Only Author and Title</u>

**Wang Y, Liu F, Liu X, Shi S, Bi Y, Moejes FW (2019) Comparative transcriptome analysis of four co-occurring Ulva species for understanding the dominance of Ulva prolifera in the Yellow Sea green tides. J Appl Phycol 31: 3303–3316**

Google Scholar: <u>Author Only Title Only Author and Title</u>

**Wells ML, Potin P, Craigie JS, Raven JA, Merchant SS, Helliwell KE, Smith AG, Camire ME, Brawley SH (2017) Algae as nutritional and functional food sources: revisiting our understanding. J Appl Phycol 29: 949–982**

Google Scholar: <u>Author Only Title Only Author and Title</u>

**Wichard T, Charrier B, Mineur F, Bothwell JH, Clerck OD, Coates JC (2015) The green seaweed Ulva: a model system to study morphogenesis. Front Plant Sci. doi: 10.3389/fpls.2015.00072**

Google Scholar: <u>Author Only Title Only Author and Title</u>

**Wichard T, Oertel W (2010) Gametogenesis and Gamete Release of Ulva Mutabilis and Ulva Lactuca (chlorophyta): Regulatory Effects and Chemical Characterization of the “Swarming Inhibitor”1. Journal of Phycology 46: 248–259**

Google Scholar: <u>Author Only Title Only Author and Title</u>

**Wu C, Jiang P, Guo Y, Liu J, Zhao J, Fu H (2017) Isolation and characterization of Ulva prolifera actin1 gene and function verification of the 5’ flanking region as a strong promoter. Bioengineered 0: 1–10**

Google Scholar: <u>Author Only Title Only Author and Title</u>

**Yamazaki T, Ichihara K, Suzuki R, Oshima K, Miyamura S, Kuwano K, Toyoda A, Suzuki Y, Sugano S, Hattori M, et al (2017) Genomic structure and evolution of the mating type locus in the green seaweed Ulva partita. Scientific Reports 7: 11679**

Google Scholar: <u>Author Only Title Only Author and Title</u>

**Zheng Z, He B, Xie X, Wang G (2020) Co-suppression in Pyropia yezoensis (Rhodophyta) reveals the role of PyLHCI in light harvesting and generation switch. Journal of Phycology. doi: 10.1111/jpy.13073**

Google Scholar: <u>Author Only Title Only Author and Title</u>

**Zhou L, Wang L, Zhang J, Cai C, He P (2016a) Complete mitochondrial genome of Ulva linza, one of the causal species of green macroalgal blooms in Yellow Sea, China. Mitochondrial DNA Part B 1: 31–33**

Google Scholar: <u>Author Only Title Only Author and Title</u>

**Zhou L, Wang L, Zhang J, Cai C, He P (2016b) Complete mitochondrial genome of Ulva prolifera, the dominant species of green macroalgal blooms in Yellow Sea, China. Mitochondrial DNA Part B 1: 76–78**

Google Scholar: <u>Author Only Title Only Author and Title</u>

